# Real Time *In vivo* Analysis of Pancreatic Beta-cell Autophagic Flux Reveals Impairment Before Onset of Autoimmune Diabetes

**DOI:** 10.1101/2023.05.23.541935

**Authors:** Olha Melnyk, Charanya Muralidharan, Bryce E. Duffett, Alissa N. Novak, Glorian Perez-Aviles, Michelle M. Martinez, Justin J. Crowder, Amelia K. Linnemann

## Abstract

The catabolic pathway of autophagy is critical for pancreatic beta-cell function and is defective in established type 1 diabetes (T1D). However, it is unclear when and how this critical cell process becomes defective during diabetes pathogenesis. To study the nature of autophagy dysfunction in the context of autoimmune diabetes, we utilized intravital microscopy to study autophagic flux *in vivo* in real time. We generated a custom AAV8- packaged mCherry-eGFP-LC3B biosensor driven by the insulin promoter for beta-cell-selective expression. For real time autophagic flux evaluation, fluorescent signal from eGFP and mCherry fluorophores was correlated in space and time to follow the process of autophagosome-lysosome fusion. We observed autophagic flux defects in the beta-cells of non-obese diabetic (NOD) mouse model of T1D prior to hyperglycemia onset that were less apparent in mice without a functional immune system. We also evaluated autophagic flux in human donor islets that were transplanted under the kidney capsule of immune incompetent mice. Collectively, we provide the first evaluation of autophagic flux *in vivo* in 4D and demonstrate that autophagy defects precede hyperglycemia in NOD mice suggesting a potential causative role for these defects in beta-cell demise during T1D pathogenesis.

## Introduction

Autophagy is a complex and dynamic process that all cells, including the insulin-producing beta-cells (*1, 2*), use to degrade and recycle damaged and aging cellular components. During the process of autophagy, cytosolic components are packaged into double-membraned vesicles, called autophagosomes, and delivered to the lysosomes for recycling and degradation (*3-5*). Functional autophagy is critical for cellular homeostasis and stress response, and defects in autophagy can lead to an accumulation of undegraded cargo and proteins, ultimately leading to apoptosis (*6, 7*). It is perhaps no surprise then, that dysfunctional autophagy has been linked to a broad array of human diseases (*8*), including autoimmune diseases (*9*). However, excessive elevations in autophagy can also lead to cell death (*10*), highlighting that autophagy must be regulated within a narrow window to enable the beneficial effects while minimizing the negative effects under normal conditions.

Type 1 diabetes (T1D) is an autoimmune disease characterized by immune attack of the insulin-producing pancreatic beta-cells, leading to their destruction and the development of clinical hyperglycemia. An emerging concept in the T1D research field is that beta-cell dysfunction may play a causative role in disease pathogenesis through the alteration of the beta-cell–immune cell interface, thus contributing to the autoimmune attack of the beta-cells (*11*). We recently observed that autophagy is disrupted in the context of human T1D, and that disposal of damaged/dysfunctional beta-cell components is perturbed in autoantibody-positive human organ donors even prior to diabetes onset (*12*). These data are suggestive that defective beta-cell autophagy may play a role in the onset of hyperglycemia, rather than merely as a consequence of it. Indeed, we also observed evidence of dysfunctional beta-cell autophagy in a mouse model of spontaneous autoimmune diabetes, the nonobese diabetic mouse (NOD), prior to the development of hyperglycemia. Further, it is well established that disruption of murine beta-cell autophagy leads to impaired beta-cell function and glucose intolerance in response to pathological stress (*13-15*).

However, despite indications that defects in autophagy may contribute to beta-cell dysfunction and autoimmune diabetes pathogenesis, a limitation in the prior work is that the use of standard approaches to study autophagy primarily relies on an evaluation of *in vivo* tissues as snapshots in time (*16*). Given that autoimmune diabetes development is a stochastic process in both humans and the NOD mouse model, this effectively provides a “moving target” when it comes to determining the relationship between autophagy disruption and beta-cell dysfunction.

To address this limitation, we used a beta-cell selective fluorescent autophagy biosensor coupled to intravital imaging to visualize beta-cell autophagic flux *in vivo* in real-time using 2- photon microscopy (*16-18*). We evaluated both baseline and stimulated autophagic flux in endogenous pancreatic islet beta-cells of the NOD mouse model of spontaneous autoimmune diabetes and in immune incompetent NOD *scid* gamma mice (NSG), as well as in human islets transplanted under the kidney capsule of NSG mice. Our data demonstrate defects in baseline and IFN-α stimulated autophagic flux in pre-diabetic NOD mice, that is likely due to a combination of genetics and immune infiltration. We are also able to monitor a heterogeneous response of human donor islets to IFN-α-induced autophagy. Importantly, IFN-α is associated with early T1D pathogenesis (*19, 20*), suggesting that this defective response in the pre-diabetic phase play a role in early dysfunction associated with disease development.

## Results

### Autophagy biosensor successfully labels LC3B protein in autophagosomal membrane within beta-cells *in vitro* and *in vivo*

The set of available fluorescent proteins to study autophagy is constantly growing (*21-24*), allowing functional assessment of autophagy in addition to more comprehensive observation of different autophagic vesicles. Here, we used a dual-fluorescent biosensor (*17, 25*) combined with real-time fluorescent microscopy (*26*) to understand the complex kinetics of autophagic flux *in vivo*.

We utilized the previously designed (*27, 28*) dual-fluorescent eGFP (*29*) and mCherry (*30*) biosensor conjugated to the microtubule-associated protein 1B-light chain 3 (LC3B). eGFP is a green fluorescent protein with enhanced stability in the cytoplasm, but is sensitive to low pH. During autolysosome formation, the fluorescent signal from eGFP is quenched due to the decreased pH level within the lysosome. However, the red mCherry fluorescent signal stays intact during the process of autophagosome binding with the lysosome (Fig 1A). The biosensor thus changes from emitting both “red” and “green” fluorescent signals when the LC3B protein is connected to the autophagosome, to carrying “red” signal only when the autophagosome is bound with the lysosome.

**Figure 1.**
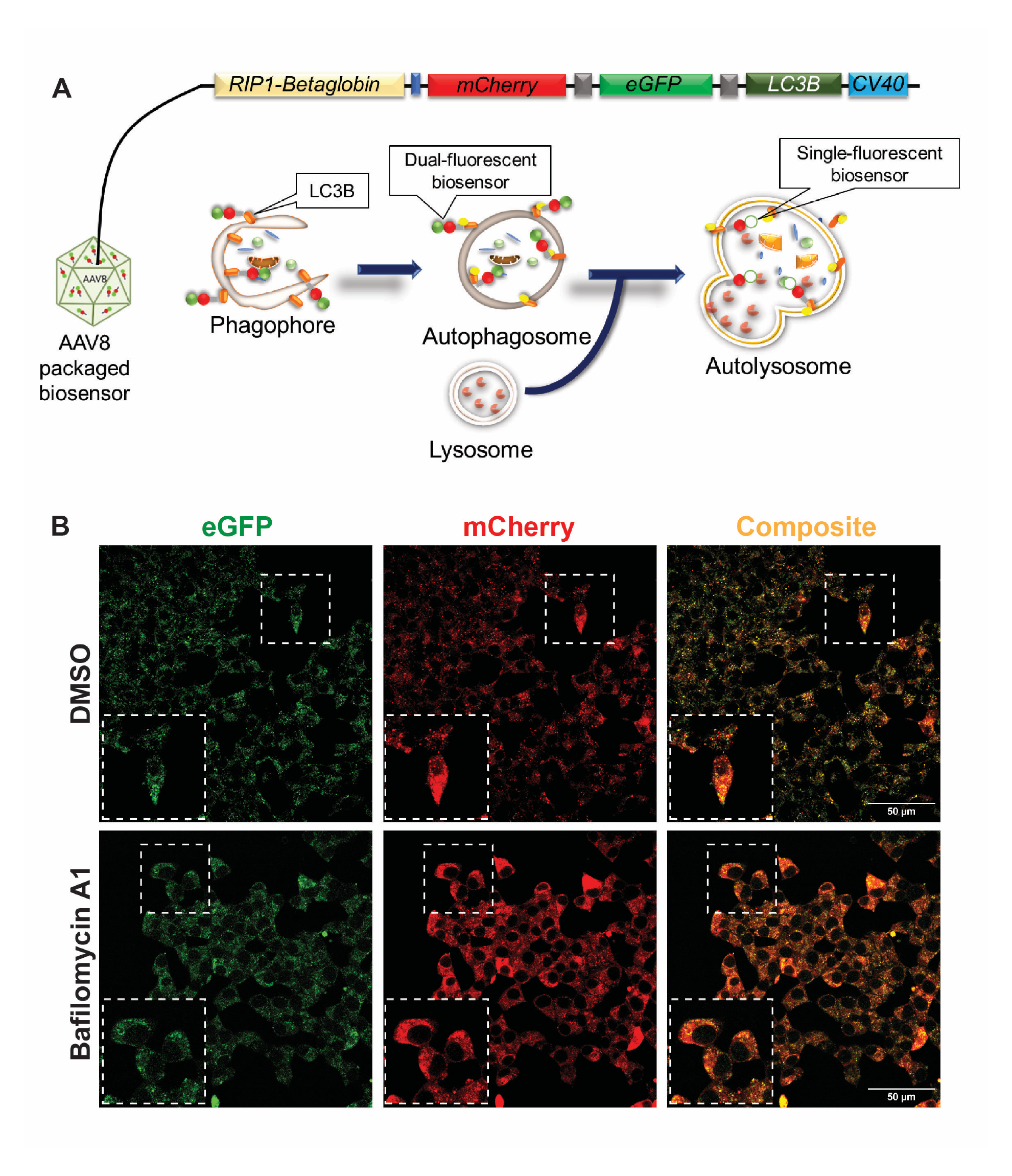
Autophagy biosensor successfully labels autophagosomal membrane and functions as expected within beta-cells *in vitro*. **(A)** Schematic showing the macroautophagy process with AAV8- LC3B-eGFP-mCherry biosensor binding site and expected fluorescence change. **(B)** Representative images of INS-1E cells transduced with 5000 MOI of autophagy biosensor then treated with the autophagy inhibitor Bafilomycin A1. The enlarged site is shown in the frame for each image. Scale bar = 50 μm.

The eGFP-mCherry-LC3B biosensor was packaged in the adeno-associated virus serotype 8 (AAV8) and delivered to the beta-cells, as described using other sensors in our previous work (*31*). We first validated the biosensor *in vitro* using rat insulinoma cells (INS-1 832/13 cells (*32*)), a model cell line for the pancreatic beta-cells (Fig 1B). Cells were transduced with the biosensor for 24 hours and then treated with either vehicle (DMSO) or Bafilomycin A1, an autophagy inhibitor (a vacuolar H^+^-ATPase inhibitor) (*33, 34*), for 1 hour to verify biosensor function. Cells were fast fixed with 4% paraformaldehyde and imaged to identify “red” and “green” puncta, representing the autophagosomes. Following the Bafilomycin A1 treatment, we observed an increase in colocalized signal compared to DMSO treated cells, indicating an accumulation of autophagosomes and demonstrating that the beta-cell-selective promoter does not impair expected biosensor function.

We proceeded to validate the biosensor *in vivo*. A cohort of C57BL/6J mice were intraperitoneally (IP) injected with the AAV8 packaged biosensor, and then intravital imaging was performed on the endogenous pancreas 3 weeks later. Robust biosensor expression was observed in several islets throughout the pancreas (Fig 2A). Following this confirmation, mice were retro-orbitally injected with saline and imaged in 4D (X+Y+Z+Time) to measure the basal level of autophagy, thus providing an internal control for each islet. The autophagic vesicle turnover events have been evaluated previously in immortalized cell lines (*35, 36*), and these data suggest that autophagic turnover occurs within approximately 10-15 min in these cells. Therefore, taking into consideration the autophagy turnover time frame, images were acquired every 2 min for 30 min, to account for one full autophagy turnover process of at least one vesicle in each cell.

**Figure 2.**
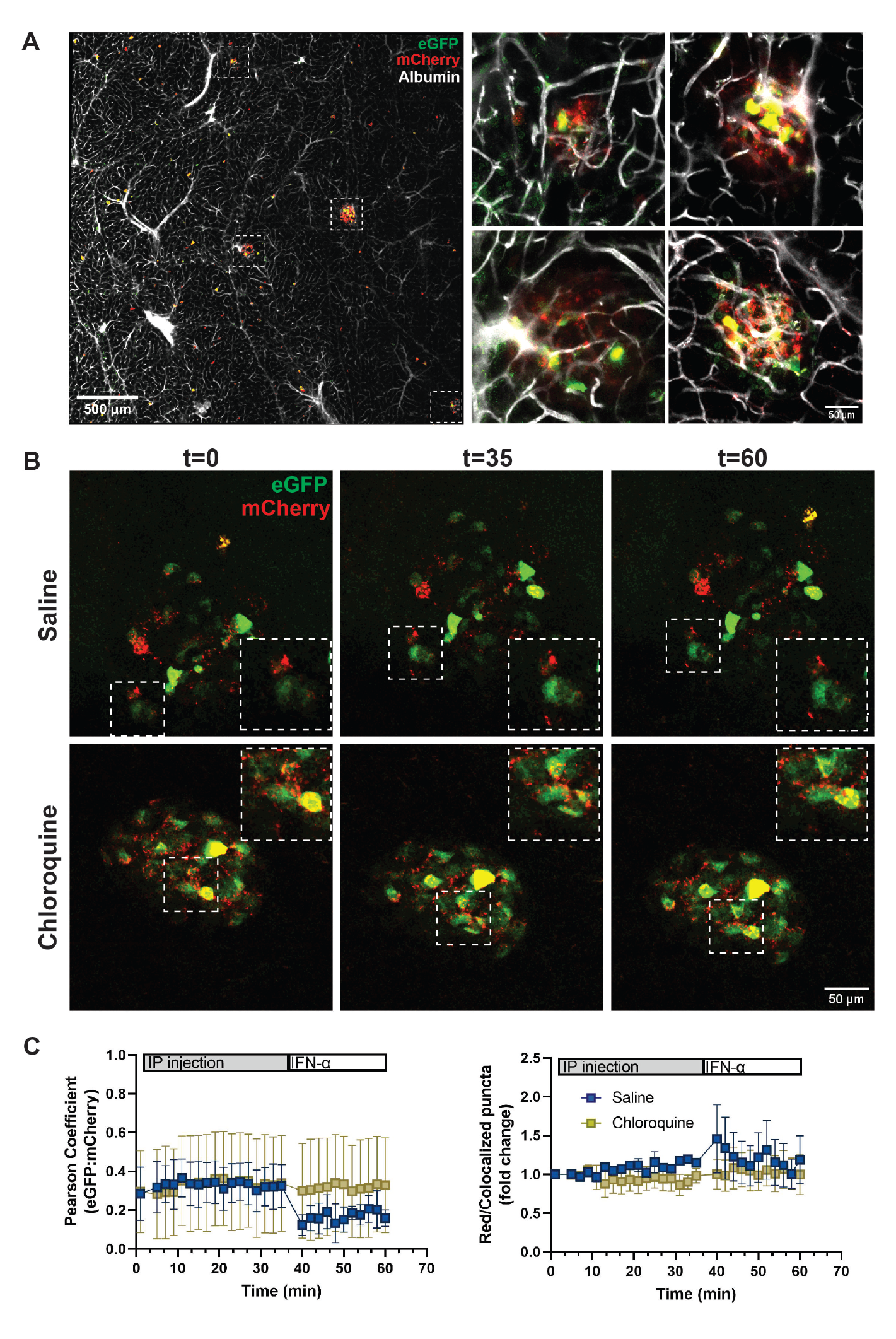
The autophagy biosensor effectively measures autophagic flux in wild-type mice. **(A)** Tiled representative image of a large area of a C57BL/6J mouse pancreas expressing the AAV8-delivered autophagy biosensor, with multiple islets identified. Identified islet are enlarged as a maximum image projection of the collected z-stacks. **(B)** Representative images of islets at various time points during intravital imaging. Selected cells are enlarged in the corner of each image. **(C)** Quantitative analysis of autophagic flux in wild-type mice. Pearson correlation coefficient and the puncta count in the form of red/colocalized puncta ratio fold change are shown for mice injected with either saline followed by IFN-α or chloroquine followed by IFN-α. Data points are mean +/- SEM, N=4 animals per condition.

We can evaluate changes in autophagic flux using compounds that stimulate or inhibit autophagy. Therefore, these same mice were then injected with recombinant mouse IFN-α and autophagic flux was continually monitored for an additional 20 minutes. IFN-α is a type I interferon, which is produced as part of the innate immune response to viral infection. Importantly, we chose this cytokine not only because type I interferons have been shown to stimulate autophagy (*37*), but also because IFN-α–mediated signaling is a key component of T1D pathophysiology. Specifically, it has been demonstrated that children genetically at risk for T1D have a type I IFN-inducible transcriptional signature in blood cells that precede the appearance of autoantibodies (*19, 20*). Representative images throughout this imaging are shown (Fig 2B), whereas videos of all images collected are shown in the supplemental materials.

For analysis of autophagic flux, two parameters were quantified. First, the Pearson correlation coefficient of “red” + “green” colocalized signal was used to assess the presence of autophagosomes over time in the beta-cells. The ratio of “red” only to colocalized (“red” + “green”) puncta was used to assess the conversion to autolysosomes, and thus, the rate of autophagosome degradation. When autophagy is stimulated, we expect a decrease in the Pearson correlation coefficient (i.e., a decrease in autophagosome number) and an increase in the red/colocalized puncta ratio (i.e., autolysosomes). When autophagy is inhibited, we expect that the Pearson coefficient and the ratio of red/colocalized puncta will remain unchanged. Using this approach, we observed that saline infusion had no effect on autophagic flux, whereas IFN-α infusion was able to rapidly stimulate autophagic flux, as indicated by a reduction in autophagosomes (“red” + “green” signal) and a corresponding increase in autolysosomes (“red” signal normalized to “red” + “green” signal; Fig. 2C). Alternatively, in the presence of the autophagy inhibitor chloroquine, the baseline presence of autophagosomes was increased and unchanged in response to IFN-α whereas the presence of autolysosomes was globally reduced and again unchanged in response to IFN-α (Fig. 2C).

### Autophagic flux is impaired in prediabetic NOD mice

Our prior data from static evaluation of prediabetic NOD mice and autoantibody positive human donors suggested that autophagic flux may be disrupted before the onset of hyperglycemia (*12*). Therefore, we turned our attention to determining whether autophagic flux is impaired in the NOD model before the mice become hyperglycemic. A series of NOD mice, and immune incompetent NSG mice as a control, were injected with the autophagy biosensor as above. Intravital imaging of the pancreas in NSG mice indicated that islets could be readily identified by biosensor expression (Fig. 3A). Pancreata were collected from the animals injected with biosensor, then embedded and sectioned to evaluate infection efficiency throughout the entire pancreas in these models. The mCherry signal from the biosensor remains throughout tissue processing and can be used to mark biosensor infected cells, whereas insulin antibody can be applied to specifically mark all beta-cells (Fig. 3B). Quantification of multiple tissue sections from across the entire pancreas revealed a large proportion of beta-cells expressing biosensor within individual islets (Fig. 3C), which was representative of 40-80% of islets throughout the pancreas expressing biosensor in at least 30% of their beta-cells (Fig. 3D). Collectively, these data demonstrate overall efficient biosensor infection of the islets in the mouse strains evaluated by intravital imaging.

**Figure 3.**
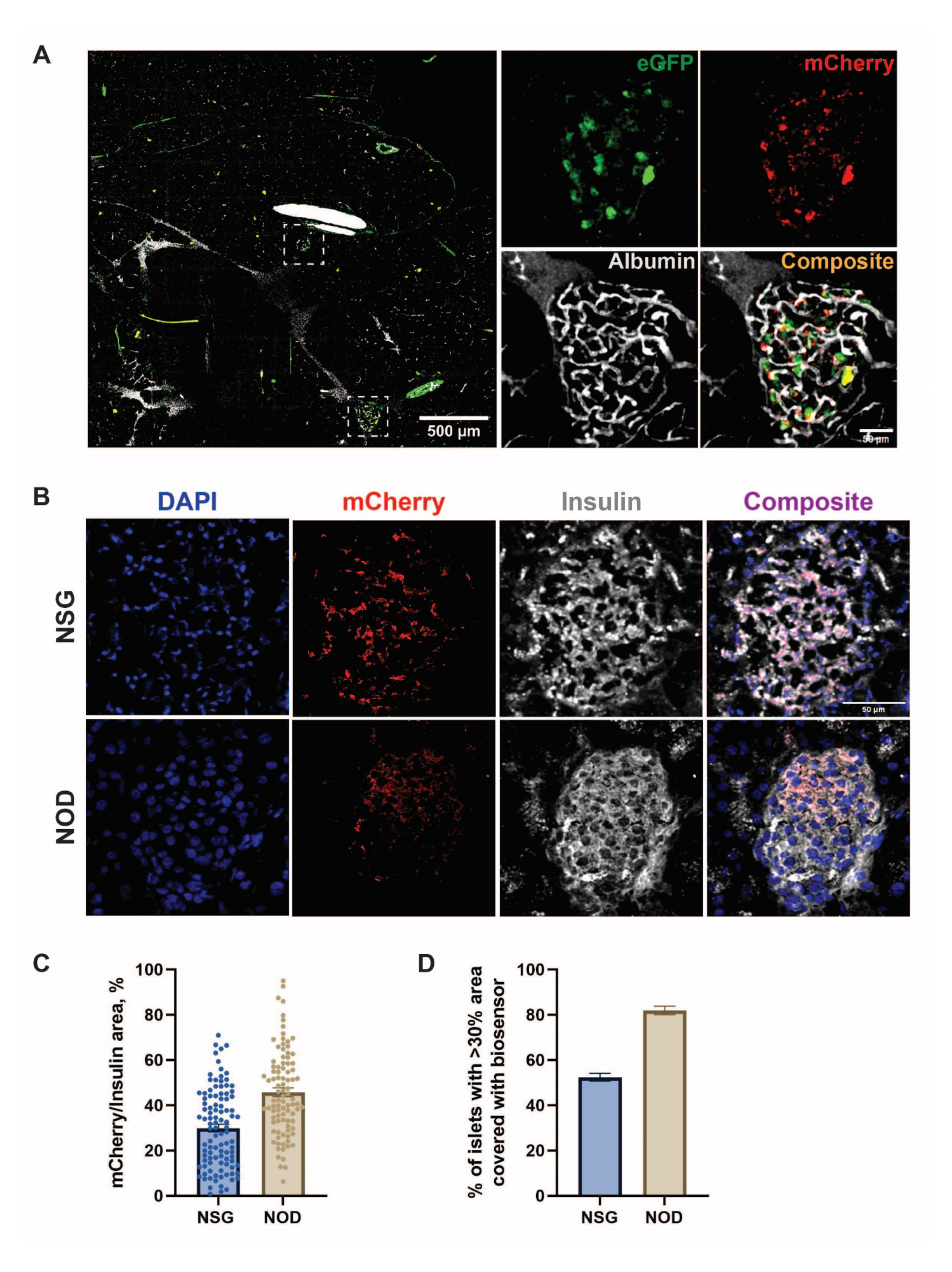
NOD and NSG beta cells efficiently express the autophagy biosensor. **(A)** Intravital image of a large area of biosensor infected NSG mouse pancreas with two individual islets denoted (white boxes). One islet is shown at higher resolution as a maximum projection of the collected z-stack. **(B)** Representative images of islets in pancreas sections from female NSG and NOD mice after intravital imaging. **(C)** Quantification of (B) showing the proportion of beta-cells within a given islet that are also expressing biosensor (i.e., the mCherry positive area). **(D)** Quantification of the percentage of the islets throughout the pancreas expressing biosensor in at least 30% of the beta-cells. N=6 mice per group.

To investigate autophagic flux in the beta-cell prior to the onset of diabetes, we proceeded to perform intravital imaging of 10-week-old NOD mice injected with the autophagy biosensor. We included both female and male NOD mice, which develop diabetes stochastically at different rates. Female NSG mice were used as genetically matched controls. Pancreatic islets expressing biosensor were identified in the endogenous pancreas as above, and baseline z-stacks of the selected islet were collected. We then injected a series of mice with saline or chloroquine followed by IFN-α, and collected images over time, as above (Fig. 4A). The Pearson correlation coefficient and puncta count analysis were then performed to assess autophagic flux in 4D (XYZT). We observed that IFN-α was able to stimulate a reduction in autophagosomes and a concomitant increase in autolysosomes in female NSG mice, though these effects were less robust than those observed in mice on the B6/J background (Fig. 4B). Notably, female NOD mice exhibited no response to IFN- α whereas male NOD mice had a reduced response relative to the NSG controls. Together, these data suggest impaired autophagic flux in the NOD genetic background that is likely further impaired in the presence of autoimmunity. This conclusion is further supported by the inhibition of response to IFN-α by chloroquine in the beta-cells of female NSG and male NOD mice, but a lack of effect in the beta-cells of female NOD mice (Fig. 4C). Overall, these results clearly demonstrate impaired autophagic flux prior to onset of hyperglycemia in the NOD model of autoimmune diabetes.

**Figure 4.**
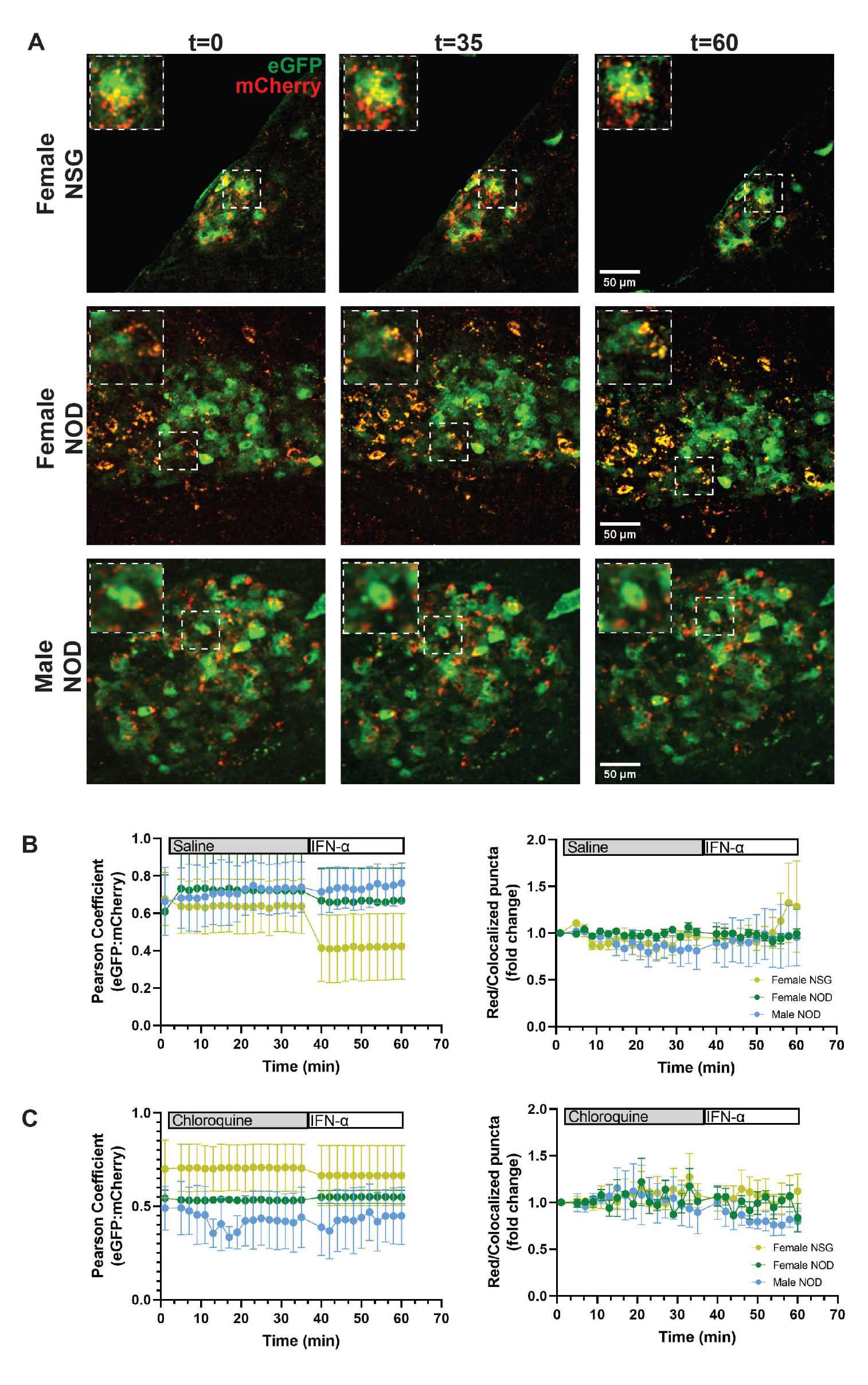
The autophagy biosensor reveals the impairment of autophagic flux of the female and male NOD mice compared to the female NSG control. **(A)** Representative images of the analyzed islets are shown for female NSG, female NOD, and male NOD mice after saline and IFN-α injections. Single cell images depicted in the frames are enlarged. **(B)** The autophagic flux quantitative analysis is shown for female NSG, female NOD, and male NOD mice after saline and IFN-α injections. Pearson correlation coefficient as a function of time and red/colocalized puncta fold change over time is shown. Mean values with SEM are plotted for each animal strain. N=3-4 for each strain, 1-2 islets per animal analyzed. **(C)** The quantitative analysis of autophagic flux after Chloroquine and IFN-α injection is shown. Pearson correlation coefficient as a function of time and red/colocalized puncta fold change over time is shown. Mean values with SEM are plotted. N=3-4 animals per strain with 1-2 islets per animal analyzed.

### Autophagic flux is heterogeneous in human islet beta-cells in an *in vivo* setting

Many limitations preclude the study of human islet autophagic flux in a true *in vivo* state within the endogenous pancreas. However, as a proxy to studying human beta-cells in their true niche, our approach to study autophagic flux *in vivo* in real-time can also be used to study human beta-cells by transplanting them into NSG mice. Although human islets can be transplanted in several areas of a mouse, where they will engraft effectively (*38*), one of the most successful and easily accessible places for human islet engraftment is the space under the kidney capsule. We therefore transduced human donor pancreatic islets *in vitro* with the autophagy biosensor, then transplanted them under the kidney capsule of NSG mice. Islets from each individual donor were transplanted into 4 mice. After two weeks, to allow for engraftment of the islets, mice were subjected to intravital imaging of the kidney containing human islets expressing biosensor (Fig. 5A).

**Figure 5.**
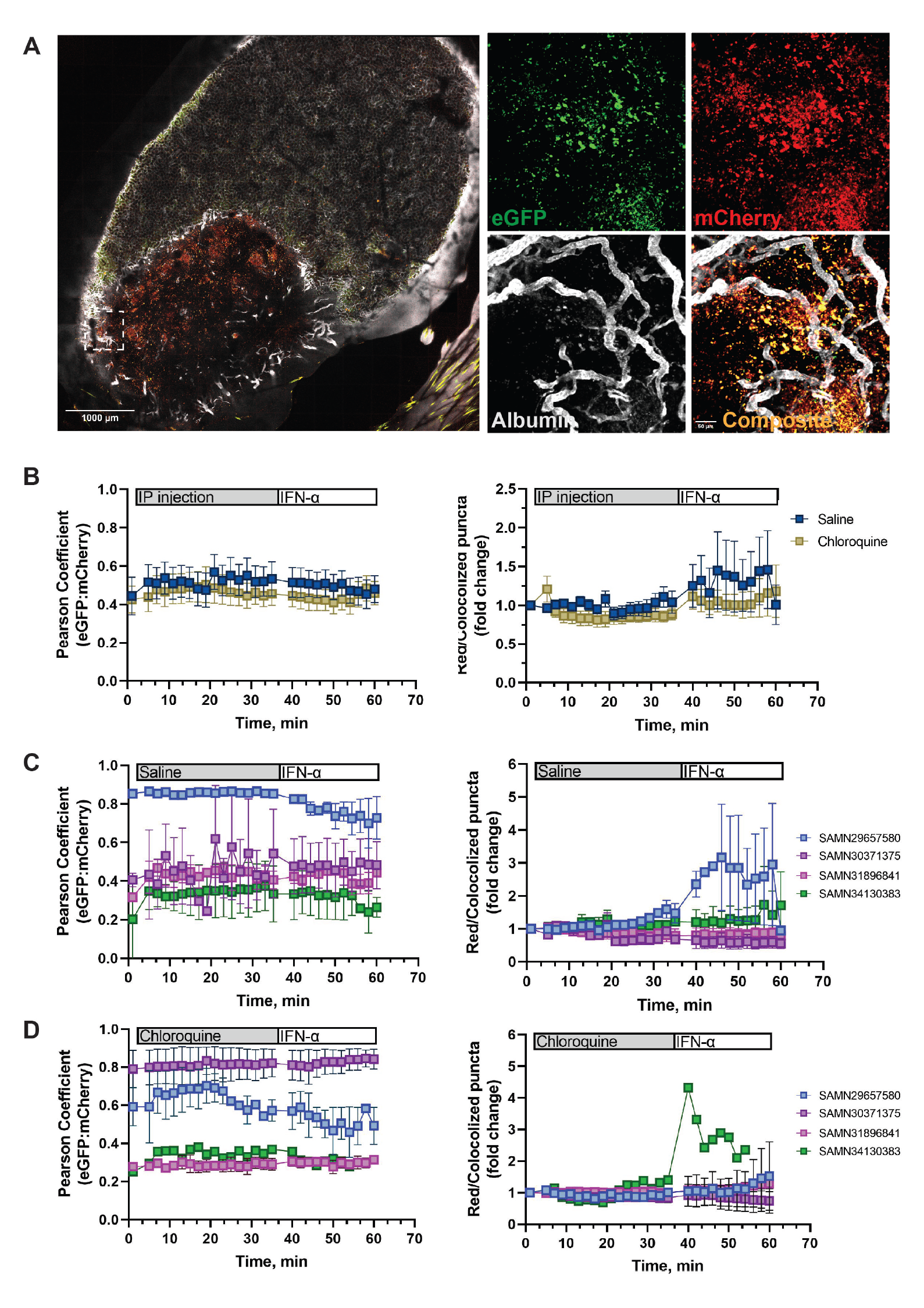
Human islet autophagic flux is heterogeneous *in vivo*. **(A)** Representative large-scale image of an NSG mouse kidney containing transplanted human islets infected with the autophagy biosensor with a region of interest (white box) further highlighted to show the EGFP (green) and mCherry (red) of the biosensor. The vascular network in and around the islets are labeled by Albumin conjugated to AlexaFluor-647 (white). **(B)** Quantitative analysis of autophagosomes (left) and autolysosomes (right) in all donor islets exposed to saline/IFN-α (blue) or chloroquine/IFN-α (gold). Mean values +/- SEM are shown for N=3 donors, with data representative of 2 mice per donor per condition. **(C)** Quantitative analysis of autophagosomes (left) and autolysosomes (right) separated by individual donor showing response to saline/IFN-α. Mean values +/- SEM are shown for each donor. **(D)** Quantitative analysis of autophagosomes (left) and autolysosomes (right) separated by individual donor showing response to chloroquine/IFN-α. Mean values +/- SEM are shown for each donor.

Using a similar treatment approach as above for mouse pancreas, we then proceeded to evaluate autophagic flux in the transplanted human islets. For each donor (Table 1), a set of animals were first injected with saline to evaluate baseline autophagic flux. We then followed this with an injection of recombinant human IFN-α to stimulate autophagy and the response was imaged in 4D (XYZT). Another parallel set of mice engrafted with islets for each donor were injected with chloroquine followed by IFN-α. When evaluating all donors together (Fig. 5B), we observed that the stimulation of autophagic flux by IFN-α injection is modest compared to the effect observed in mice. Chloroquine abolished these modest effects (Fig. 5B).

**Table 1.**
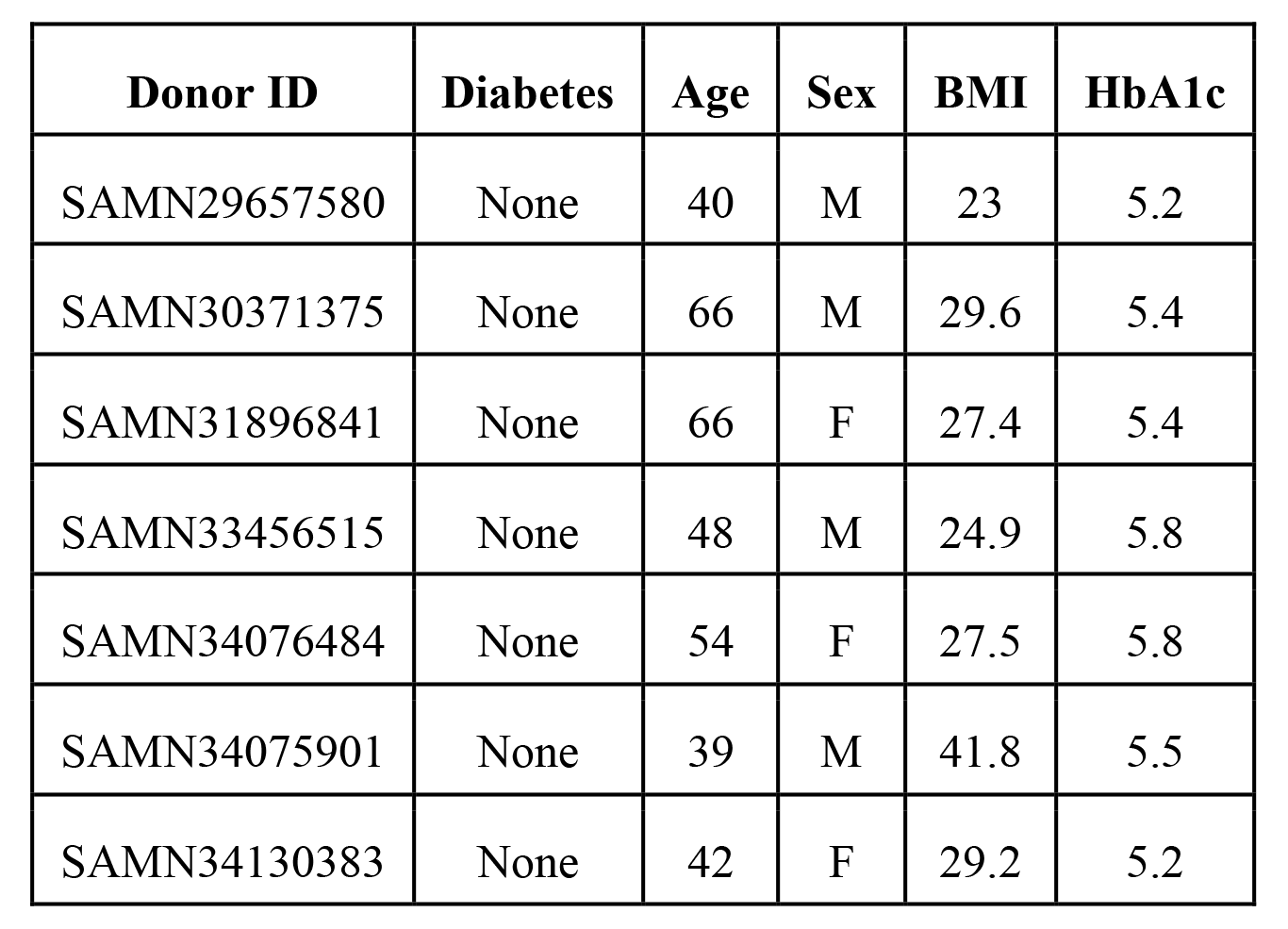
Human organ donor information.

| **Donor ID** | **Diabetes** | **Age** | **Sex** | **BMI** | **HbA1c** |
| --- | --- | --- | --- | --- | --- |
| SAMN29657580 | None | 40 | M | 23 | 5.2 |
| SAMN30371375 | None | 66 | M | 29.6 | 5.4 |
| SAMN31896841 | None | 66 | F | 27.4 | 5.4 |
| SAMN33456515 | None | 48 | M | 24.9 | 5.8 |
| SAMN34076484 | None | 54 | F | 27.5 | 5.8 |
| SAMN34075901 | None | 39 | M | 41.8 | 5.5 |
| SAMN34130383 | None | 42 | F | 29.2 | 5.2 |

It is well established that interindividual variability between human donors can impact upon islet physiology and function (*39*). Therefore, we also evaluated autophagic flux in each donor individually. Interestingly, when individual donor data was plotted (Fig. 5C and D), we observed a heterogenous response to IFN-α that was consistently abolished by the presence of chloroquine for all donors. *In vitro* analysis of human islet autophagy stimulation by IFN-α confirms a heterogeneous response between donors and suggests that the *in vivo* heterogeneity is likely not due to engraftment efficiency (Fig. S1). These data collectively demonstrate that similar to mouse pancreatic islets, autophagic flux can be stimulated in human islets by IFN-α exposure and that this can be imaged in an *in vivo* setting in real time. However, further investigation will be required to identify the factors responsible for the heterogeneous autophagic response of human donor beta-cells to IFN-α stimulation.

## Discussion

T1D has classically been viewed solely as an immune driven disease. However, emerging evidence suggests that beta-cells may play an active role in their own demise (*40-42*). Autophagy plays an important role in beta-cell homeostasis and survival (*1*). In fact, it has been extensively studied in the context of type 2 diabetes, beta-cell homeostasis (*2*), and ER stress response (*43*). However, autophagy has only recently been investigated in the context of T1D (*44*).

Multiple approaches can be used to study autophagy such as collection and preservation of tissues for further investigation with established techniques of immunofluorescence, western blotting, and electron microscopy (*16, 17, 45*). However, limitations exist in that many of these approaches provide only a static snapshot of the dynamic autophagy process. Virally packaged fluorescent biosensors are a useful and versatile alternative to study dynamic processes, and we have previously used this approach to evaluate reactive oxygen species dynamics and calcium oscillations in beta-cells *in vivo* with great success (*31, 38, 46, 47*). Fluorescent autophagy sensors have also been used successfully to study autophagic flux in beta-cells *in vitro* by us and others (*48, 49*). Therefore, we reasoned that this approach could be used to enable the investigation of the dynamic nature of autophagy and allow the visualization of autophagic flux *in vivo*.

Autophagy sensors have been used to evaluate autophagy *in vivo* using isolated tissues from transgenic mice (*50, 51*), and while studies such as this provide useful information, they stop short of truly evaluating real time dynamic autophagic flux in the presence of systemic modulators. Here, we have coupled the use of a validated beta-cell-selective dual-fluorescent biosensor (Fig. 1) with intravital imaging to interrogate the dynamics of autophagic flux in real-time *in vivo.* The use of viral packaging of the sensor allows us to study beta-cell autophagy in any mouse model or in isolated islets without complicated breeding to genetically introduce the sensor, or the risk of negative consequences from prolonged expression of multiple fluorescent proteins but with high infection efficiency (Fig. 3). We therefore used this versatile approach to selectively study beta-cell autophagic flux *in vivo* in both endogenous mouse islets and in xenografted human islets.

To study autophagy dynamically in the context of T1D, we used a mouse model of spontaneous autoimmune diabetes (NOD mice). Using both male and female mice, we evaluated beta-cell autophagy at different stages of the immune attack on pancreatic islets prior to diabetes onset in the NOD mice. We chose to inject animals with the cytokine IFN-α because of its established relevance to human T1D pathogenesis (*19, 20*), with the hypothesis that it should stimulate autophagic flux as part of the innate response to infection, but only under conditions where autophagy is able to be stimulated normally. Indeed, our testing in wild type C57B6/J mice demonstrated a clear stimulation of beta-cell autophagy by IFN-α (Fig. 2). However, we observed an impairment in autophagic flux by IFN-α in pre-diabetic female NOD mice compared to the age-and sex-matched NSG control (Fig. 4). This provides a clear demonstration that autophagy is impaired prior to the onset of diabetes in the NOD model, consistent with our previous observations using NOD tissues *ex vivo*, and our observations suggesting defects in autophagy in autoantibody positive human donors (*12*).

While the above establishes that autophagy is defective prior to diabetes onset in the NOD model, the question as to the impact of autoimmunity on autophagy suppression remained. In other words, our data thus far was unable to fully distinguish between the effects of genetic predisposition and immune mediated repression of beta-cell autophagy. Therefore, we evaluated autophagic flux in age-matched male NOD mice. It is well known that male NOD mice develop diabetes much later and with a lower penetrance than their female counterparts (*52*), so the local immune cell infiltration should be reduced compared to females at the same age. We indeed observed increased stimulation of autophagic flux by IFN-α in the beta-cells of male NOD mice compared to female NOD mice (Fig. 4). However, the ability to stimulate autophagy in these animals was still somewhat repressed compared to the NSG and C57B6/J mice. We interpret this to mean that there is likely a combination of genetics and autoimmunity that contribute to autophagy suppression in the NOD model. However, we also acknowledge that direct causality cannot be adequately judged using these models, and additional studies must be carried out to definitively answer this question. Given that we have now established the platform for 4D observation of autophagic flux in the endogenous pancreas *in vivo* and have established that autophagic flux is impaired prior to hyperglycemia, these and other future studies to establish the role of autophagy in the natural history of autoimmune diabetes are now a reality.

Although human islets cannot be studied *in vivo* in the same way as mouse islets for ethical reasons, human islets are widely studied in an *in vivo* context after transplantation into NSG mice (*38*). Therefore, we proceeded to use this approach to perform real-time 4D autophagy observation in human beta-cells in intact islets *in vivo* after transplantation under the kidney capsule of NSG mice. Using recombinant human IFN-α, we observed that while we could sometimes stimulate autophagic flux, there exists a significant amount of donor heterogeneity in autophagy in the nondiabetic donors evaluated (Fig. 5). This is perhaps not surprising and may be reflective of interindividual biological variability that is not replicated in an inbred mouse colony, as we have used. However, islet isolation can induce quite a bit of stress that may persist even after transplantation, and it has been reported that autophagic activity can be impaired by metabolic stress in a heterogeneous manner (*51, 53*). Nonetheless, we demonstrate an ability to stimulate human islet autophagic flux in human islets *in vivo*, thus establishing our platform for future studies in islets from donors with diabetes and/or for testing therapeutic autophagy modulators.

In summary, we demonstrate, for the first time in 4D, that beta-cell autophagic flux is impaired prior to hyperglycemia in an established model of autoimmune diabetes. Further, we can monitor human beta-cell autophagy in 4D in an *in vivo* setting. Collectively, our observations not only open a new line of questioning for the mechanisms controlling autophagy loss in the pathogenesis of T1D, but provide a platform for evaluating the efficacy of novel drugs that could be used to modulate this critical process. The potential for studies such as these to evaluate not only acute drug response, but also long-term response and functional outcomes through the incorporation of abdominal imaging windows (*31, 54, 55*) remains to be seen. However, we anticipate that many new discoveries will emanate from this work in future studies to contribute to our understanding of T1D pathogenesis.

## Materials and Methods

### Mice

C57BL/6J (RRID:IMSR_JAX:000664) and non-obese diabetic (NOD; RRID:IMSR_JAX:001976) mice were purchased from The Jackson Laboratory (ME, USA) at 6 weeks of age. NOD.Cg-Prkdc*^scid^* Il2rg*^tm1Wjl^*/SzJ (NSG; RRID:IMSR_JAX:005557) mice were obtained from the on-site breeding colony at the Preclinical Modeling and Therapeutics Core (PMTC) at Indiana University Simon Comprehensive Cancer Center at 6 weeks of age. Mice were housed in a temperature-controlled facility with a 12 h light/12 h dark cycle and were given free access to food and water. All experiments were approved by the Indiana University School of Medicine Institutional Animal Care and Use Committee. Blood glucose for NOD mice was monitored bi-weekly with an AlphaTrak2 glucometer (Zoetis, NJ, USA), and were characterized as diabetic after 2 consecutive days of blood glucose readings >250 mg/dL. Male and female NOD, and female NSG mice at the age of 7 weeks were injected with ∼2.31×10^12^ GC of the AAV8 biosensor and intravital imaging was performed at the age of 10 weeks.

### Generation of a beta-cell-selective fluorescent autophagy biosensor

AAV8 has previously been shown to have pancreas tropism (*56*), and we have previously used a hybrid insulin/rabbit beta globin promoter (*57*), which we refer to as RIP1, to enable beta-cell-selective sensor expression *in vivo* (*31*). Here, we generated AAV8-RIP1-mCherry-EGFP-LC3B plasmid by cloning the insulin/beta globin promoter and the mCherry-EGFP-LC3B construct (from Addgene construct #22418, (*28*)) into an AAV8 packaging plasmid. The plasmid was then packaged into the adeno-associated virus serotype 8 (AAV8) at the Vector Core of the Gene Therapy Program at the University of Pennsylvania.

### Transduction of endogenous mouse pancreatic islets

To induce beta-cell-selective expression of the RIP1-mCherry-EGFP-LC3B biosensor, animals were given intraperitoneal injections of AAV8-packaged sensor three weeks prior to the intravital imaging session. All injections were given to animals at 7 weeks of age and between 9am and 11am to ensure consistency of sensor expression in the tissue. This enables robust expression of sensor specifically in the pancreatic beta-cells, that will last for at least 20 weeks beyond the injection time (Fig. S2).

### Transduction and transplantation of human islets

Human islets were obtained via the Integrated Islet Distribution Program (IIDP) or from the Alberta Diabetes Institute (ADI) islet core. Islets were cultured in standard Prodo islet media (Prodo, PIM-S001GMP; 5.8 mM Glucose) containing 5% human AB serum supplement (Prodo, PIM-ABS001GMP), 1% Glutamine and Glutathione supplement (Prodo, PIM-G001GMP), and 10mg/mL Ciproflaxacin (Fisher, MT61277RG). Upon receipt, the islets were centrifuged at 350g for 5 min to remove shipping media and then gently resuspended in complete islet media to recover overnight before infection with AAV8-packaged biosensor. After overnight recovery of the islets, the islets were centrifuged at 350g for 5 min at room temperature. The supernatant was aspirated, and the pelleted islets were incubated with Accutase (Fisher Scientific) for 1 minute at 37°C. This leads to a “loosening” of the islets, but not a full dissociation of the cells, and aids in increasing infection efficiency of the biosensor. The enzyme was inactivated with islet media containing serum and then aspirated after centrifugation. Islets were then transduced with 5000 MOI of AAV8-RIP1- mCherry-EGFP-LC3B for ∼16-18hrs at 37°C incubator in complete islet media.

Following transduction, the islets were centrifuged at 350g for 5 min and viral islet media was removed. The islets were then transferred to virus-free islet media and transplanted under the left kidney capsule of NSG mice. Approximately, 1000 IEQs were transplanted per mouse. Briefly, mice receiving the transplants were subcutaneously treated with Ethiqa XR (3.25mg/kg) prior to starting the surgery. During the surgery, mice were anesthetized through isoflurane inhalation and kept on a warming blanket until they regained reflexes post-surgery. The left flank of the mouse (around the kidney area) was shaved, and excess hair was removed with Nair hair removal cream (Church & Dwight Co., Inc.). Following, the shaved area was cleaned with an alcohol swab and a small incision on the skin was created. The left kidney was gently externalized. The distal end of a sterile polyethylene (PE-50) tubing was attached to a Hamilton screw syringe containing a blunt needle tip. The tubing containing the islets was cut at an angle to make the tubing beveled. Then, a small nick was made on the kidney capsule with a 30-gauge needle and the beveled end of the tubing containing the islets was carefully placed under the kidney capsule, all while keeping the tissue moist with saline. Then a small air bubble was delivered very carefully and slowly under the capsule followed by islets. The kidney was then placed back into the peritoneal cavity and the muscle and skin were sutured using 5-0 VICRYL® sutures. Post-surgery, the animals were treated with analgesics/anti-inflammatories and were monitored until they regained reflexes. The animals were singly housed in a clean cage and provided with wet feed and were subcutaneously administered Carprofen (5 μg/kg) as necessary for pain management post-surgery.

### Intravital imaging of endogenous mouse pancreas

Mice were anesthetized by isoflurane inhalation, then a small flank incision was made and pancreata were gently externalized as we have previously described (*31*). Animals were placed such that the externalized pancreas could rest on the surface of a glass bottom imaging dish (0.17 mm thick, 40 mm in diameter, WillCo). The temperature and humidity of the externalized organ were monitored throughout the imaging session. Throughout the surgery and imaging, ophthalmic ointment was applied to avoid drying of the animal’s eyes. To reach the deeper pancreas tissue, 2- photon microscopy was utilized, with penetration depth reaching from 500 µm up to 1 mm into the tissue (*58, 59*). Animals were imaged using a Leica TCS SP8 DIVE confocal multiphoton system mounted on an inverted stand equipped with a Spectra-Physics MaiTai DeepSee laser and mounted with a 40x 1.1 NA water objective. Intraperitoneal injection (IP) and retroorbital injections were performed on the animals during the imaging. Initially pancreatic islets were identified through the eyepiece by looking for the “red” and “green” fluorescence of the beta-cells expressing the autophagy biosensor. We collected 512×512 pixel images at 400 Hz scanning frequency with 900 nm and 1050 nm wavelength excitation lasers, whereas emission signal from the biosensor was collected on the Power HyD detectors at 490-530 nm and 575-625 nm emission windows for eGFP and mCherry signals, respectively. Next, a baseline 4D z-stack of each selected islet was collected (2-3 µm thick) non-stop for a duration of 3 min. Following this, animals were IP injected with 100 μl saline or chloroquine diphosphate (50 mg/kg; Tocris Bioscience) and selected islet was imaged for a total duration of 30 min with images collected at time intervals of 2 min. Animals were then given retroorbital injections of 2.75×10^5^ IU of recombinant mouse IFN-α (BioLegend), and the final 20 min of images with the same time intervals were collected. To evaluate islet architecture, retroorbital injection of albumin conjugated with Alexa Fluorophore 647 was performed at the end of saline/chloroquine/IFN-α imaging. Images of the vasculature were collected using an 840 nm laser wavelength, with emission collected in the 650-700 nm wavelength window of the PMT detector.

### Intravital imaging of human islets

A similar procedure as the intravital imaging for mouse pancreas was performed for the human islet imaging. During the surgery before the imaging session, mice under inhaled isoflurane anesthesia underwent gentle externalization of the left kidney, which was placed on the glass-bottomed imaging dish. The kidney was kept moist throughout all procedures with saline and the temperature of the imaging dish was kept near body temperature levels (36-37°C). Next, each animal was placed on the microscope and imaged using a 25x 0.95NA water objective. Transplanted human islets were identified first through the eyepiece and then a z-stack of 2-3 μm thick slices was imaged for 3 min non-stop at 400 Hz frequency to obtain a baseline sensor readout.

IP injection of the saline or chloroquine was performed as above and then chosen islet was imaged for 30 min at 2 min time intervals. Next, retroorbital injection of recombinant human IFN-α (2.75×10^5^ U; Novus Biologicals) was performed and the last 20 min of images of the same islet at the same time interval was collected.

### Analysis of intravital images

Images were collected in sequential order: the first excitation of 900 nm laser was set, and appropriate emission spectra were collected, the next 1050 nm laser excitation was set. The images were collected sequentially between laser lines, but fast switching allowed overall image collection to be performed almost simultaneously. FIJI was used for all analysis (*60*). The maximum intensity projection of the collected z-stacks of the islet was created first for further image analysis. Collected image channels from 900 nm laser excitation and 490-530 nm emission and 1050 nm laser excitation and 575-625 nm emission were then separated and analyzed for colocalization and puncta count. As an initial step, background subtraction was performed on both images: image background was identified in the places of images where no signal is visible, and the average intensity of background pixels was subtracted from the whole image. Next, rolling ball background subtraction was performed with the FIJI plugin, of radius 25 pixels. For the red and green colocalization in space, the Pearson correlation coefficient was measured for the selected images using the colocalization threshold plugin in ImageJ. The Pearson correlation coefficient for the positive area was used as a measured parameter. Next, a median filter was applied to the images for puncta identification in both channels. Using the Image calculator plugin in FIJI, the AND image of red and green channels was created (only pixels which have positive intensity simultaneously in both channels will be shown) and identified as a colocalized signal. The Particles Analysis plugin in ImageJ was then used to calculate puncta. To avoid puncta overcounting, a percentage of the positive area was used as a measuring parameter for the puncta count. The Red Only image was created as a difference between colocalized and red signal images, to count only red puncta which do not have green signal associated with them. The same Particle Analysis plugin was used to count the Red Only puncta. Next ratio of the Red Only/Colocalized puncta was plotted for each time data point. Finally, data were represented as a fold change of the time 0 data point (baseline image collected for 3 min prior to any injections done to the animals).

Pearson correlation coefficient is insensitive to the intensity difference between two channels, thus we only analyze cooccurrence in space of the two fluorophores. Puncta count specifically evaluates the autophagic flux. With the increase of the autophagic flux, the number of red puncta will increase, and yellow will decrease simultaneously, leading to an increased ratio. In the case of autophagy inhibition with chloroquine, the number of red puncta will not change significantly, but the number of yellow puncta will increase, leading to a decreased ratio.

### Immunofluorescence staining of mouse islets after intravital imaging

After imaging, pancreata were collected and flash frozen in Optimal Cutting Temperature (OCT) compound. OCT compound embedded pancreata were sectioned at 8 µm thickness, then the tissues were permeabilized with 1% PBS containing Triton x100 for 15 min. Slides were then blocked with Dako blocking buffer (Agilent Technologies) prior to incubation with antibodies in Dako antibody diluent (Agilent Technologies) overnight at 4°C in a humidified chamber with the following primary antibodies: Guinea-pig anti-Insulin antibody (Biorad 5330-0104G, 1:250). Following overnight incubation, the slides were washed with PBS, and secondary antibodies Goat anti-Guinea pig Alexa Fluor 647 (Invitrogen) and Donkey anti-Rabbit Alexa Fluor 488 (Invitrogen) were added at 1:500 dilution together with DAPI at a 1:2000 dilution to stain nuclei. The slides were incubated in the dark with secondary antibodies for 1 hour at room temperature and then mounted using a hard-set mounting medium (Everbrite). All images were acquired using Zeiss LSM 700 confocal microscope, with 40x 1.3NA oil objective. Collected 10 µm thick z-stacks (with 1µm step) of the islets images from the tissue slices were projected as a maximum intensity on one plane then background subtraction was performed (by measuring the mean intensity of the not fluorescent part of the image and subtracting that value from whole image) for all channels separately, and an insulin mask was created to measure the insulin area. The insulin mask was then applied over the mCherry, channels and the area of each signal was calculated over the insulin mask. Data was represented as a ratio of each channel area over the insulin-positive area.

## Supporting information

B6J Saline Video

B6J Chloroquine Video

Female NOD Saline Video

Female NOD Chloroquine Video

Female NSG Saline Video

Female NSG Chloroquine Video

Male NOD Saline Video

Male NOD Chloroquine Video

## Acknowledgments

The authors acknowledge the support of the Microscopy Core of the Indiana Diabetes Research Center (P30DK097512).

## Author contributions

Methodology: CM, OM, AKL

Investigation: OM, CM, BED, ANN, GPA, MMM, JJC

Supervision: AKL

Writing—original draft: OM, AKL

Writing—review & editing: OM, CM, BED, ANN, GPA, MMM, JJC, AKL

