## Supplemental Figures for "Real Time *In vivo* Analysis of Pancreatic Beta-cell Autophagic Flux Reveals Impairment Before Onset of Autoimmune Diabetes"

**Supplementary Materials**

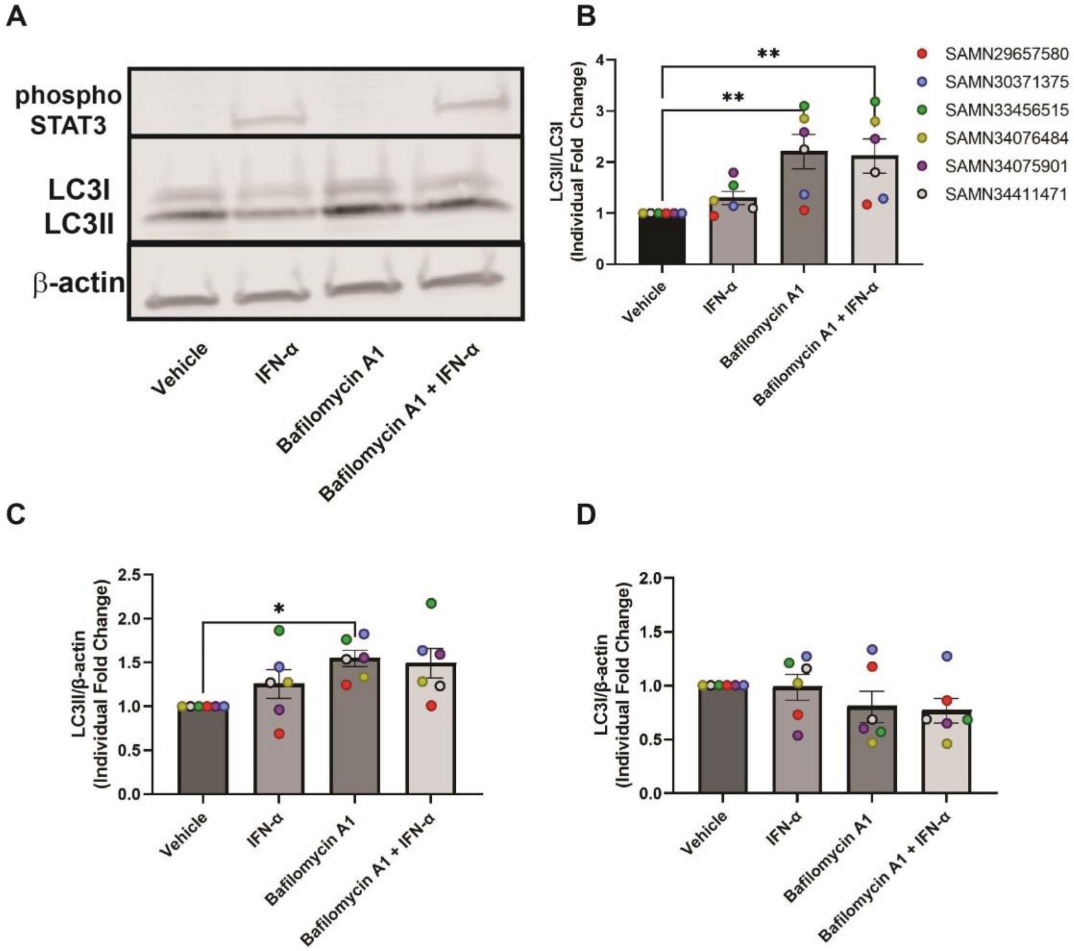


**Figure S1.** ***In vitro*** **assessment of human islet autophagic flux.** Islets in 5.8 mM glucose media were treated with 100 nM Bafilomycin A1 or vehicle for 30 min, and then 2000 IU/mL recombinant human IFN-α or vehicle was added for 1 hour. Islet media was then collected and high glucose media (16 mM) was added, then islets were incubated for an additional 1.5 hours before lysis of islets for analysis of LC3. **(A)** Representative western blots; **(B)** Quantification of LC3II/LC3I; **(C)** Quantification of LC3II; **(D)** Quantification of LC3I; N=6 donors. Nonparametric One-way ANOVA multiple comparison results shown. $^*p<0.05$, $^{**}p<0.01$

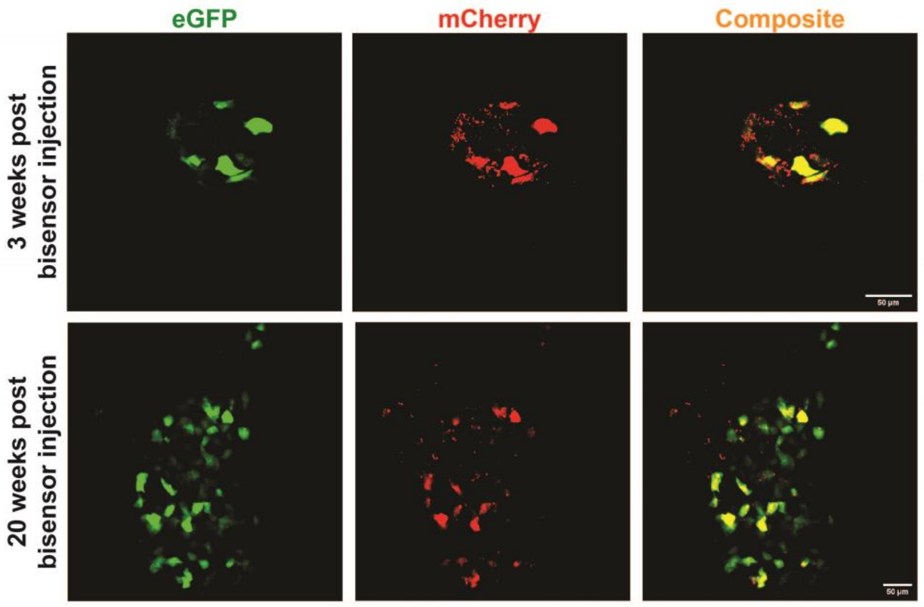


**Figure S2. Longevity of autophagy biosensor expression *in vivo*.** Representative intravital images of C57BL/6J mouse pancreatic islets expressing the AAV8-delivered autophagy biosensor at 3- and 20-weeks post biosensor injection. The EGFP (green), mCherry (red) and combined sensor expression is shown. Scale bar = 50 µm.
